# Clinicopathological characteristics and survival of lymphoepithelial carcinoma of the oral cavity and pharynx: a population-based study

**DOI:** 10.1101/669671

**Authors:** Jinbo Bai, Fen Zhao, Shuang Pan

**Affiliations:** Department of Oral and Maxillofacial Surgery, Shandong Provincial Hospital, Shandong University, Jinan, Shandong, Republic of China, 250021; Department of Stomatology, Qilu Children’s Hospital, Shandong University, Jinan, Shandong, Republic of China, 250022; Department of Orthodontics, Jinan Stomatological Hospital, Jinan, Shandong, Republic of China, 250001

**Keywords:** Lymphoepithelial carcinoma, The oral cavity and pharynx, Outcomes, SEER database, Nomogram

## Abstract

Lymphoepithelial carcinoma (LEC) of the oral cavity and pharynx is uncommon, and the characteristics and survival remains unclear. The present study aims to describe the clinicopathological characteristics and determine the factors associated with survival of this uncommon cancer. A population-based study was carried out to investigate clinical characteristics and prognosis of LEC of the oral cavity and pharynx using the data from Surveillance, Epidemiology and End Results (SEER) database between 1988 and 2013. The propensity-matched analysis was conducted for prognostic analysis, and a prognostic nomogram was also constructed. Totally, 1025 patients with LEC of the oral cavity and pharynx were identified, including 769 nasopharyngeal LEC patients and 256 non-nasopharyngeal LEC patients. The median OS of all LEC patients was 232.0m (95% CI 169.0-258.0). The 1-, 5-, 10- and 20-year survival rates were 92.9%, 72.9%, 59.3%, and 46.8%, respectively. Surgery could significantly prolong the survival time of LEC patients (P<0.01, mOS: 190m vs. 255m). Radiotherapy, as well as radiotherapy after surgery, could prolong the mOS (P<0.01 for both). The survival analysis demonstrated that old age (>60 years), lymph node (N3) and distant metastases were independent factors for poor survival, whereas radiotherapy and surgery were independent factors for favorable survival. No significant differences in survival time between nasopharyngeal LEC and non-nasopharyngeal LEC patients were observed. The prognostic nomogram was established base on five independent prognostic factors (C-index=0.70; 95% CI 0.66-0.74). In conclusion, LEC of the oral cavity and pharynx is a rare disease, and old age, lymph node and distant metastases, surgery and radiotherapy were significantly associated with prognosis. The prognostic nomogram could be used to make individual predictions of OS.

## Introduction

Lymphoepithelial carcinoma (LEC) is a rare malignant tumor which is defined as a carcinoma composed of undifferentiated malignant epithelial cells surrounded or infiltrated by a prominent component of characteristic lymphocytes and plasma cells; LEC accounts for approximately 5% of head and neck cancers[1] [2]. LEC was first described in the nasopharynx in 1921, and is also a tumor mostly located in the nasopharynx, where it represents 40% of all neoplasms[3]. Aside from the nasopharynx, this disease can also occur in other locations, including the oral cavity, salivary glands, parotid glands, etc. A previous study reported that the incidence of salivary LEC is second to nasopharynx LEC, but salivary LEC is exceedingly rare and comprises only 0.4% of salivary cancers[4].

The etiopathogenesis of LEC is not fully clear, tobacco smoking and alcohol consumption are identified to be contributing factors for its development. Nasopharynx LEC is almost invariably associated with Epstein-Barr virus (EBV) infection, and human papillomavirus (HPV) has also been identified to link to LEC of the larynx, hypopharynx and oropharynx[5, 6]. Previous studies demonstrated LEC has significant lymphocytic infiltration that could cause a strong immune response, thus, LEC patients were always accompanied with good prognosis [7, 8]. LEC is also believed to be more radiosensitive, and radiotherapy as the single modality of treatment for locoregional LEC has been described. The combination of surgery and postoperative radiotherapy has also been recommended for some non-nasopharyngeal LEC patients[7].

However, knowledge of LEC is currently limited to case reports and small case series, especially non-nasopharyngeal LEC. The clinicopathological characteristics and survival of nasopharyngeal LEC and non-nasopharyngeal LEC of the oral cavity and pharynx have not been well defined. Therefore, we performed a retrospective analysis of patients with LEC of the oral cavity and pharynx registered in SEER database to present the clinicopathological characteristics and prognosis. The characteristics and prognoses of nasopharyngeal LEC and non-nasopharyngeal LEC were also compared. Moreover, we constructed a prognostic nomogram to help physicians make an individualized survival prediction.

## Methods and Materials

### Participants

All patients with a diagnosis of LEC (ICD-O-3:8310/3, ICD-0-3/WHO 2008) located in the oral cavity and pharynx between 1988 and 2013 were identified from the SEER database. The inclusion criteria: patients with primary LEC as their only cancer. The demographic information and clinicopathological characteristics of these patients were extracted using SEER*Stat 8.2.2 software including age at diagnosis, sex, race, primary site, pathological grade, SEER historic stage classification, TNM stage, and the use of surgery, radiation and chemotherapy. The overall survival information was also identified and extracted from SEER database[9, 10]. The Institutional Review Board of Jinan Stomatological Hospital approved this study.

### Statistical analysis

Continuous data were compared using Student’s t-tests, while categorical data were examined using chi-square tests. To compare the differences in survival time between nasopharyngeal LEC and non-nasopharyngeal LEC patients, we conducted the propensity-matching (PSM) analysis with a 1:1 ratio based on age, race, pathological grade, TNM stage, and the use of surgery, radiotherapy and chemotherapy. The Kaplan-Meier method and log-rank test was used to evaluate the influence of each variable on survival time. Univariate and multivariate Cox regression survival analysis were utilized to assess the association of each variable with prognosis. The independent prognostic factors in the multivariate Cox analysis were also included to construct the prognostic nomogram. All statistical analysis was performed using MedCalc software (version 15.2.2, Mariakerke, Belgium) and R 3.1.3 software (http://www.r-project.org). P<0.05 was considered as statistically significant.

## Results

### Patients’ characteristics

Totally, 1025 patients with LEC of the oral cavity and pharynx were identified. **Table 1** depicts the characteristics of these patients and their treatment regimens. The lesions of most patients (769/1025) were located in the nasopharynx, while the non-nasopharynx LECs were observed in the salivary gland (108/1025), tonsil (79/1025), tongue (38/1025), and other sites (31/1025).

**Table 1:**
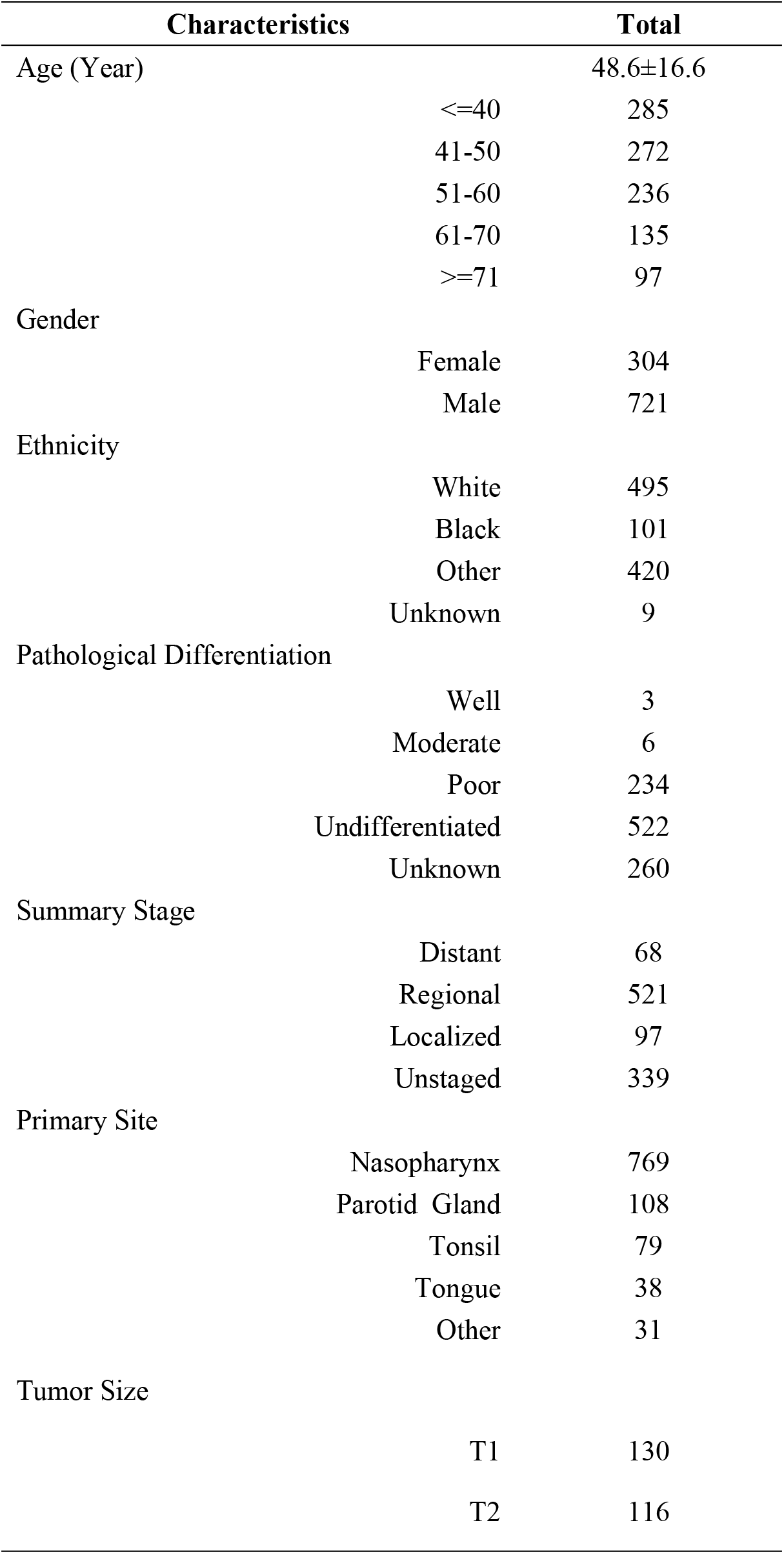
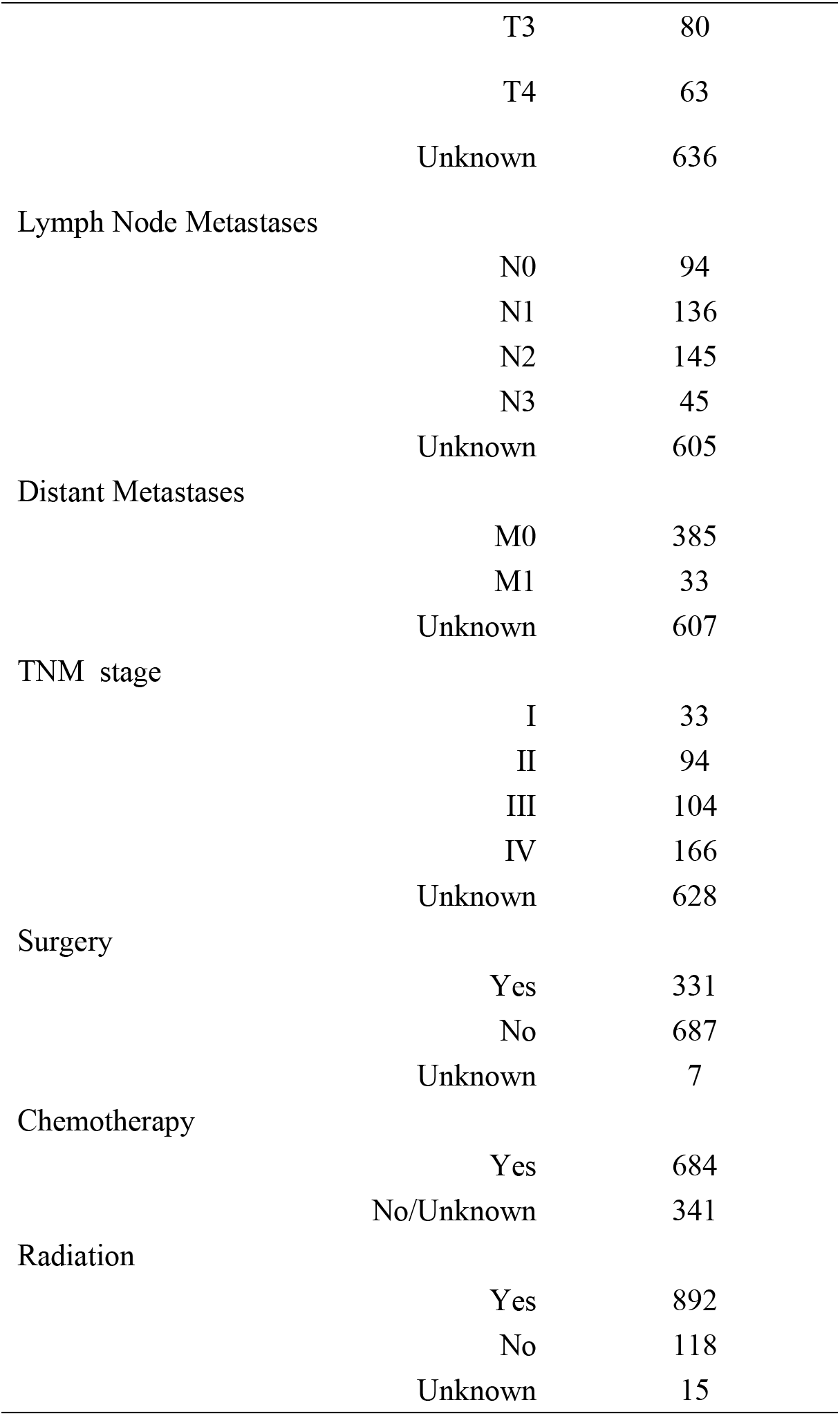
Characteristics of the 1025 patients with lymphoepithelial carcinoma of the oral cavity and pharynx

| Characteristics | Total |
| --- | --- |
| Age (Year) | 48.6±16.6 |
| ≤40 | 285 |
| 41-50 | 272 |
| 51-60 | 236 |
| 61-70 | 135 |
| ≥71 | 97 |
| Gender |  |
| Female | 304 |
| Male | 721 |
| Ethnicity |  |
| White | 495 |
| Black | 101 |
| Other | 420 |
| Unknown | 9 |
| Pathological Differentiation |  |
| Well | 3 |
| Moderate | 6 |
| Poor | 234 |
| Undifferentiated | 522 |
| Unknown | 260 |
| Summary Stage |  |
| Distant | 68 |
| Regional | 521 |
| Localized | 97 |
| Unstaged | 339 |
| Primary Site |  |
| Nasopharynx | 769 |
| Parotid Gland | 108 |
| Tonsil | 79 |
| Tongue | 38 |
| Other | 31 |
| Tumor Size |  |
| T1 | 130 |
| T2 | 116 |

|  |  |  |
| --- | --- | --- |
|  | T3 | 80 |
|  | T4 | 63 |
|  | Unknown | 636 |
| Lymph Node Metastases |  |  |
|  | N0 | 94 |
|  | N1 | 136 |
|  | N2 | 145 |
|  | N3 | 45 |
|  | Unknown | 605 |
| Distant Metastases |  |  |
|  | M0 | 385 |
|  | M1 | 33 |
|  | Unknown | 607 |
| TNM stage |  |  |
|  | I | 33 |
|  | II | 94 |
|  | III | 104 |
|  | IV | 166 |
|  | Unknown | 628 |
| Surgery |  |  |
|  | Yes | 331 |
|  | No | 687 |
|  | Unknown | 7 |
| Chemotherapy |  |  |
|  | Yes | 684 |
|  | No/Unknown | 341 |
| Radiation |  |  |
|  | Yes | 892 |
|  | No | 118 |
|  | Unknown | 15 |

### Patient survival

The median OS (mOS) of LEC patients was 232.0 months (95% CI 169.0-258.0, **Fig 1A**). The 1-, 5-, 10- and 20-year survival rates were 92.9%, 72.9%, 59.3%, and 46.8%, respectively. LEC patients with stage IV had the poorest prognosis, with a 5-year OS rate of 61.0%, compared with 92.3% for stage I, 84.6% for stage II, and 78.0% for stage III LEC patients (**Fig 1B**). Similarly, LEC patients with distant stages had significantly worse prognoses compared to individuals with localized or regional stages according to the SEER historic stage classification (P<0.01 for both); patients diagnosed with distant stage had a 5-year OS rate of 58.7%, 74.0% for localized stage and 80.0% for regional stage patients, respectively (**Fig 1C**). The prognoses of LEC patients became much worse with increasing age, increasing tumor stage and lymph node invasion (P<0.01 for all, **Figs 2A, 2B** and **2C**). Similarly, The mOS of LEC patients with distant metastases was significantly shorter than those without distant metastases (P<0.01). The mOS of LEC patients with distant metastases was only 30 months (95% CI 12.0-43.0) (**Fig 2D**). LEC patients who were black or other ethnicities also had shorter survival time than white patients (P<0.05 for both). Besides, no significant association of other variables and survival could be observed.

**Fig 1:**
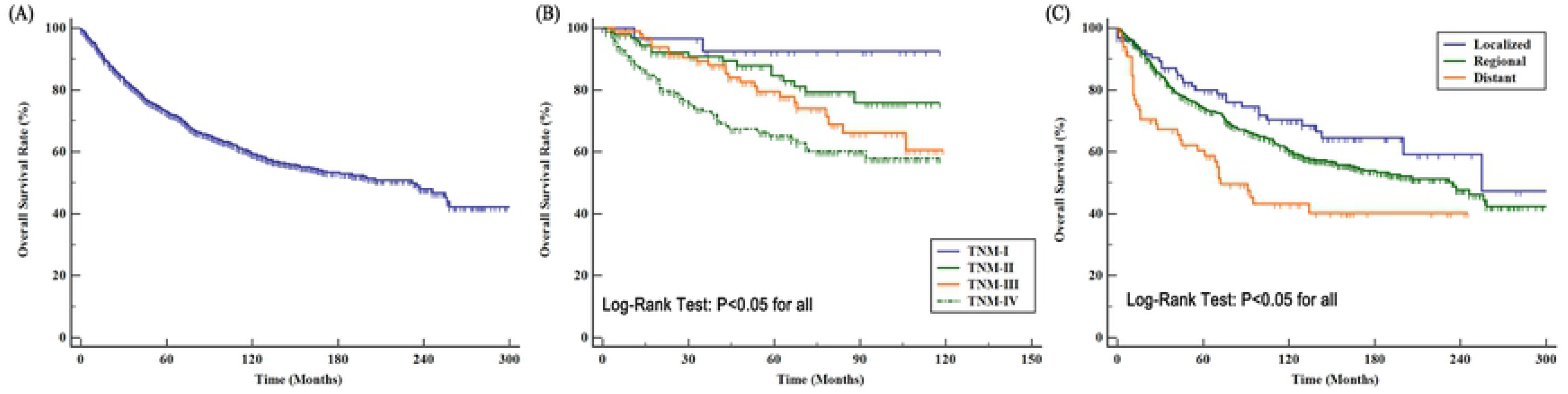
Overall survival for patients with LEC of oral cavity and pharynx; (A) overall survival for 1205 patients with LEC of oral cavity and pharynx; (B) overall survival for LEC patients with different TNM stages; (D) overall survival for LEC patients with different SEER historic stage.

**Fig 2:**
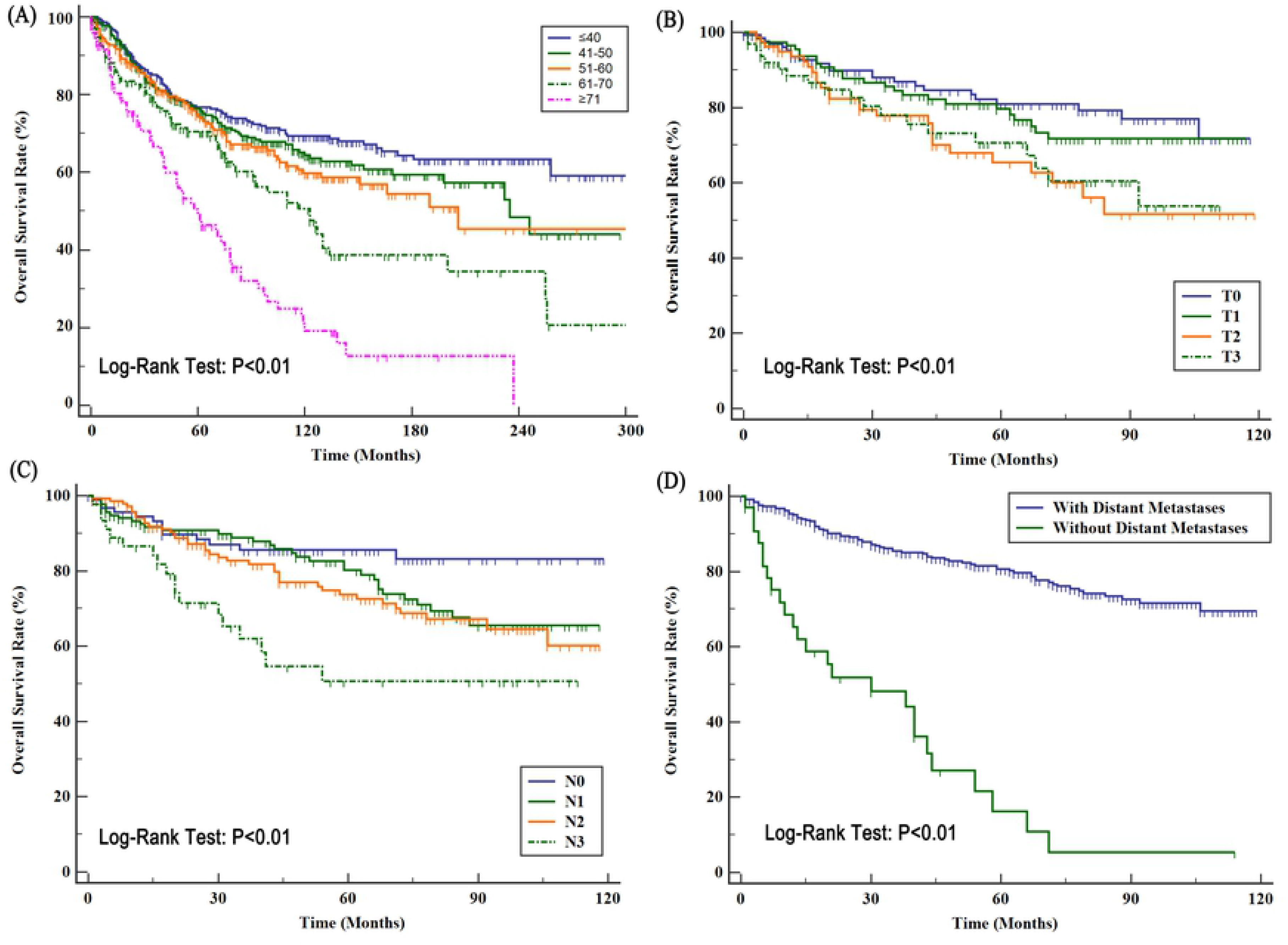
Overall survival for patients with LEC of oral cavity and pharynx; (A) overall survival stratified by age; (B) tumor stage; (C) lymph node metastases; (D) distant metastases.

### Effect of different treatments on prognosis

As seen in **Fig 3A**, surgery could prolong significantly the mOS of LEC patients (255 months vs. 190 months; P<0.01). LEC patients who received radiotherapy also had longer survival time than those without radiotherapy (P<0.01, **Fig 3B**). The mOS of LEC patients without radiotherapy was only 76 months (95% CI 43.0-138.0). However, no significant association between chemotherapy and overall survival could be observed (P=0.27). A total of 18 LEC patients received radiotherapy prior to surgery, while 239 patients received radiotherapy after surgery. The survival analysis showed that the combination of radiotherapy with surgery could significantly improve the patients’ prognoses compared with those with surgery alone (P<0.01) (**Fig 3C**).

**Fig 3:**
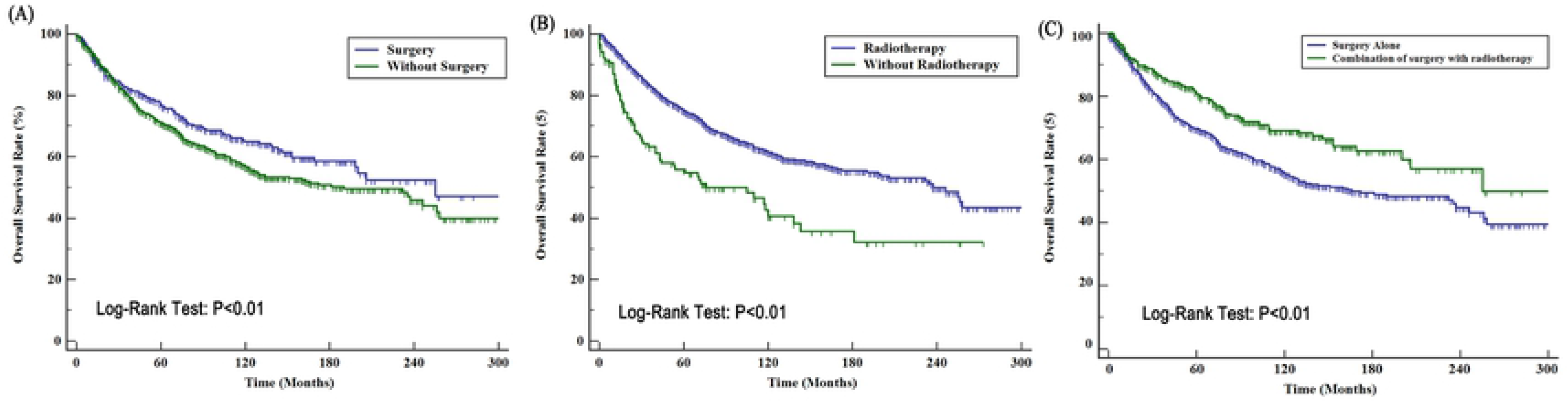
The effect of surgery and radiotherapy on overall survival for LEC patients; (A) surgery; (B) radiotherapy; and (C) combination of radiotherapy with surgery.

### Univariate and multivariate Cox proportional hazard analyses

**Table 2** showed the variables that could potentially influence OS using univariate Cox survival analysis. Old age, black ethnicity, large tumor stage, lymph node and distant metastases, and late TNM stages were significantly associated with poor prognosis, while the use of radiotherapy and surgery was related to good prognosis (P<0.05 for all, **Table 2**). Subsequently, the multivariate Cox survival analysis demonstrated that only old age (>60 years), lymph node (N3) and distant metastases were independent factors for poor prognosis, whereas radiotherapy and surgery were independent factors for favorable survival in LEC patients (**Table 2**).

**Table 2:** Univariate and multivariate Cox proportional hazard analyses of the clinical characteristics for overall survival rates in patients with lymphoepithelial carcinoma of the oral cavity and pharynx

| Factor | Category | Univariate |  | Multivariate |  |
| --- | --- | --- | --- | --- | --- |
|  |  | HR<br>(95% CI) | p-value | HR<br>(95% CI) | p-value |
| Age | <=40 | Reference |  | Reference |  |
|  | 41-50 | **1.24 (0.92-1.66)** | 0.16 | 1.01 (0.53-1.94) | 0.96 |
|  | 51-60 | **1.40 (1.04-1.90)** | 0.03 | 0.90 (0.46-1.94) | 0.75 |
|  | 61-70 | **2.03 (1.47-2.82)** | <0.01 | **1.91 (1.01-3.68)** | **0.04** |
|  | >=71 | **3.80 (2.74-5.29)** | <0.01 | **4.49 (2.11-9.56)** | **<0.01** |
| Gender | Female | Reference |  |  |  |
|  | Male | 1.19 (0.95-1.49) | 0.13 |  |  |
| Race | White | Reference |  |  |  |
|  | Black | **1.56 (1.14-2.14)** | <0.01 |  |  |
|  | Other | 1.05 (0.85-1.30) | 0.66 |  |  |
| Pathological Differentiation | Poor | Reference |  |  |  |
|  | Undifferentiated | 1.08 (0.84-1.38) | 0.55 |  |  |
| Summary Stage | Localized | Reference |  |  |  |
|  | Regional | 1.33 (0.91-1.95) | 0.14 |  |  |
|  | Distant | **2.30 (1.41-3.73)** | <0.01 |  |  |
| Primary Site | Nasopharynx | Reference |  |  |  |
|  | Non-nasopharynx | 0.85 (0.66-1.18) | 0.17 |  |  |
|  | Parotid Gland | 0.84 (0.55-1.17) | 0.25 |  |  |
|  | Tonsil | 0.69 (0.44-1.08) | 0.11 |  |  |
|  | Tongue | 0.75 (0.40-1.41) | 0.37 |  |  |
|  | Other | 1.30 (0.83-2.04) | 0.25 |  |  |
| Tumor | T1 | Reference |  |  |  |
|  | T2 | **1.23 (0.70-2.16)** | 0.47 | 1.39 (0.78-2.48) | 0.27 |
|  | T3 | **2.08 (1.19-3.63)** | 0.01 | 1.79 (0.96-3.34) | 0.06 |
|  | T4 | **2.02 (1.11-3.71)** | 0.02 | 1.39 (0.70-2.75) | 0.34 |
| Lymph Node<br>Metastases | N0 | Reference |  |  |  |
|  | N1 | **1.64 (0.87-3.12)** | 0.13 | 1.18 (0.57-2.42) | 0.66 |
|  | N2 | **1.90 (1.15-3.55)** | 0.04 | 1.67 (0.85-3.30) | 0.14 |
|  | N3 | **3.72 (1.83-7.56)** | <0.01 | **3.63 (1.56-8.48)** | **<0.01** |
| Distant<br>metastases | Yes/No | **6.98 (4.45-11.0)** | <0.01 | **6.02 (3.16-11.5)** | **<0.01** |
| Surgery | No/Yes | **0.80 (0.65-0.79)** | <0.01 | **0.65 (0.37-0.99)** | **0.04** |
| Radiation | No/Yes | **0.51 (0.36-0.74)** | 0.02 | **0.76 (0.41-0.92)** | **0.03** |
| Chemotherapy | Yes/No+Unknown | 0.89 (0.72-1.10) | 0.27 |  |  |

### Prognostic nomogram for LEC patients

To make an individualized survival prediction of LEC patients, we established a prognostic nomogram using all independent prognostic factors from the multivariate Cox survival analysis (**Fig 4**). The nomogram illustrated that the M category had the largest effect on survival, followed by age and N category. Tumor stage, race and the use of surgery and radiotherapy showed a moderate effect on prognosis. The calibration plots for the probability of overall survival at 3, 5 or 10 years in the LEC patient cohort yielded an optimal consistency between the prediction survival and the actual observation. The C-index for survival prediction in this prognostic nomogram was 0.70 (95% CI 0.66-0.74).

**Fig 4:**
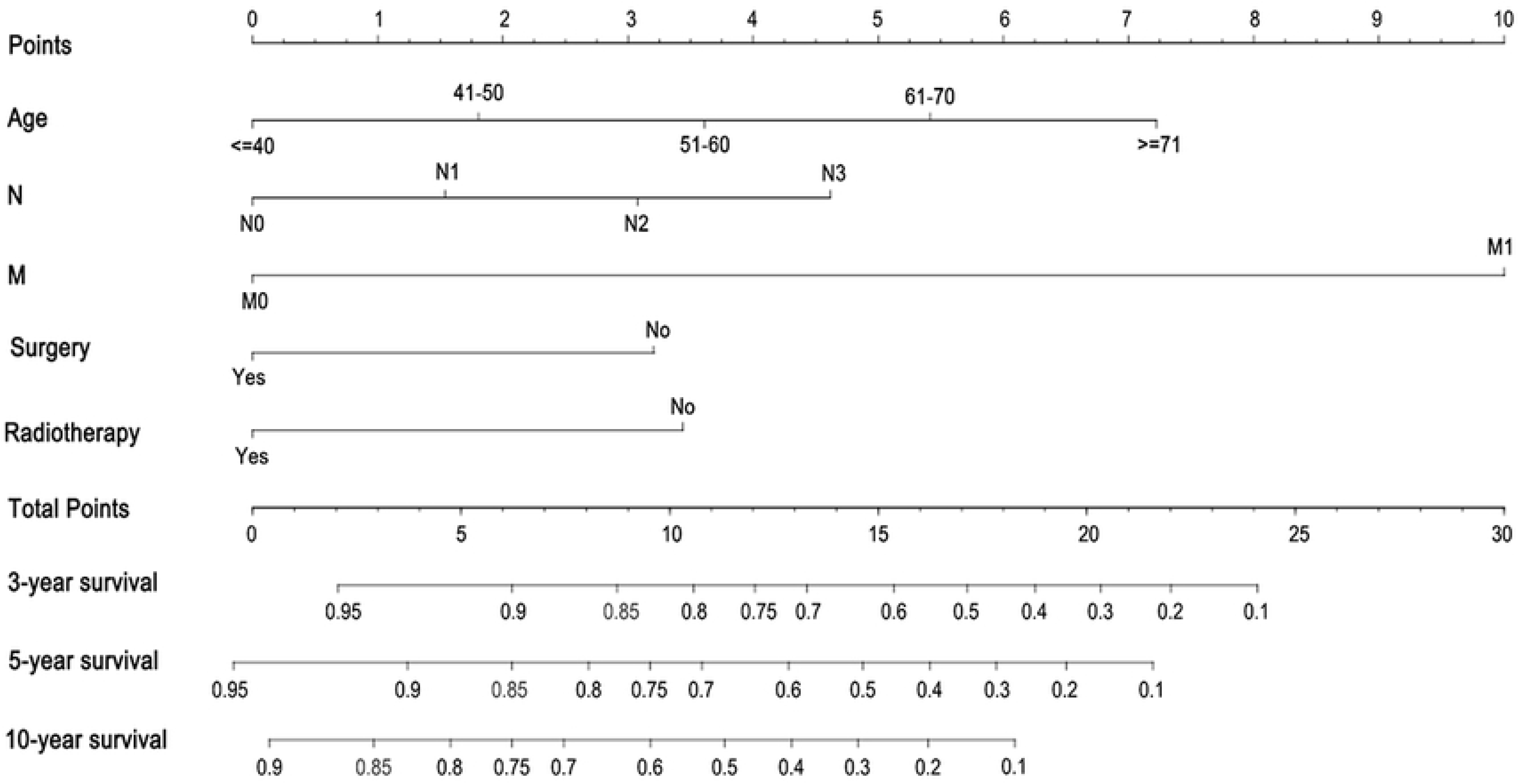
Prognostic Nomogram calculated by clinical characteristics for 3-years, 5-years, 10-years survival in patients with LEC of oral cavity and pharynx.

### Comparative analysis of nasopharynx LEC and non-nasopharynx LEC

**S1 Table** depicts the characteristics of the nasopharynx LEC and non-nasopharynx LEC patients. Non-nasopharynx LEC patients were more likely to be young and male and have an undifferentiated grade and late TNM stage than nasopharynx LEC patients, while nasopharynx LEC patients were more likely to be of American Indian/AK Native/Asian/Pacific Islander descent than non-nasopharynx LEC patients. For treatment options, non-nasopharynx LEC patients were more likely to receive surgery, radiation and chemotherapy than nasopharynx LEC patients. In the survival analysis stratified by primary site, the mOS of nasopharynx LEC patients was 235 months (95% CI 160.0-258.0), which was longer than that of non-nasopharynx patients (mOS=200m, 95% CI 143.0-256.0), but without statistically significant difference (P=0.17, **Fig 5A**). Then, we carried out a PSM analysis to balance the characteristic and treatment regimens between these two groups. A total of 96 non-nasopharynx LEC patients were matched with 96 nasopharynx LEC patients (1:1) after PSM analysis (**S2 Table**). The survival analysis demonstrated that the survival of non-nasopharynx LEC patients was slightly better than that of nasopharynx LEC patients, but without statistically significance (166.0m vs 90.0m, P=0.20, **Fig 5B**). Both the univariate Cox analysis and multivariate Cox analysis revealed that location was not significantly associated with the prognosis of LEC patients.

**Fig 5:**
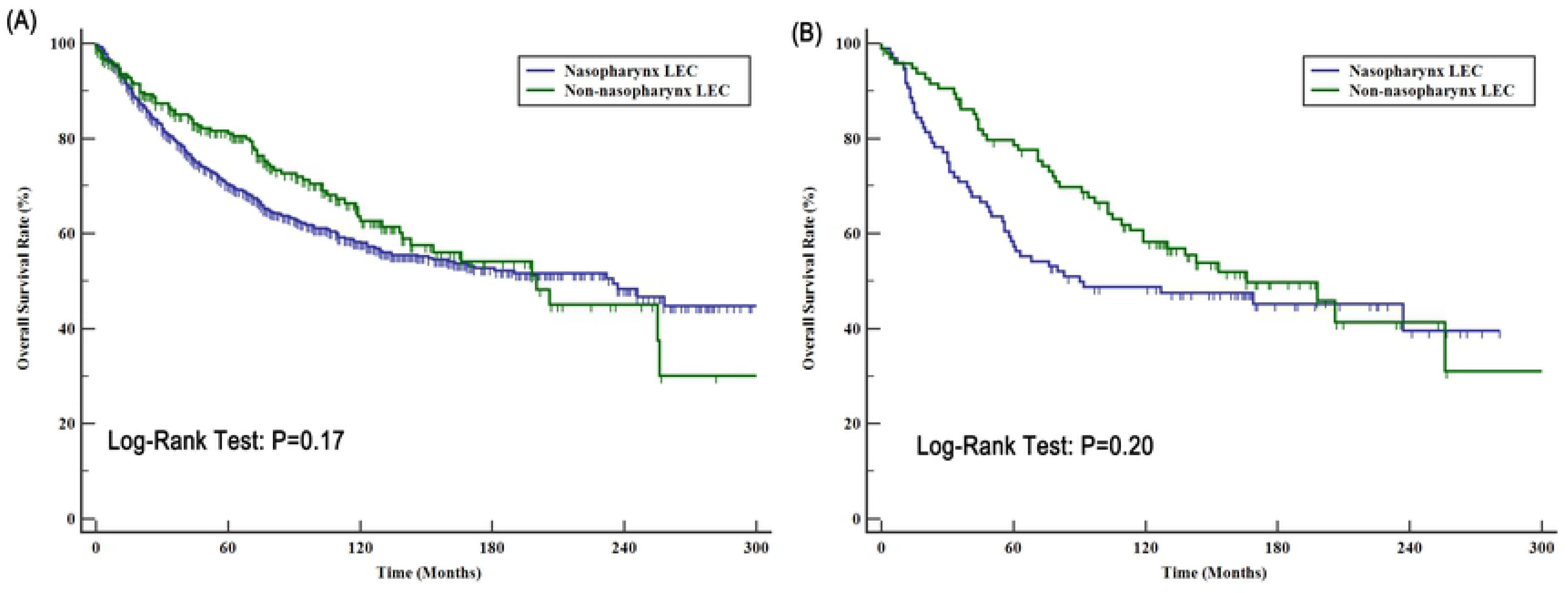
Comparative analysis of OS for nasopharynx LEC patients and non-nasopharynx LEC patients; (A) in unmatched cohort and (B) in matched cohort.

## Discussion

The majority of lymphoepithelial carcinoma studies comprises case reports or small series due to the rarity of this condition, especially non-nasopharynx LEC such as salivary gland LEC, tonsil ELC, laryngeal LEC, etc[11–13]. Therefore, the clinicopathological characteristics and survival for this disease are not fully clear, especially regarding the differences between nasopharynx LEC and non-nasopharynx LEC. In the present study, we present the clinicopathological characteristics and prognosis of those patients and determine the factors that affect survival based on the data of 1205 patients from the SEER database. We also conducted a comparison analysis of patients with nasopharynx LEC and non-nasopharynx LEC. Furthermore, we constructed a prognostic nomogram for patients with LEC of the oral cavity and pharynx to make an individualized survival prediction.

Nasopharyngeal LEC is the most commonly found in the oral cavity and pharynx. Our data showed that the tumor lesions of 75.0% of LEC patients were located in nasopharyngeal area. The patients with nasopharyngeal LEC are young, with an average age of 45.8 years. Previous studies have reported several pediatric patients with nasopharyngeal carcinoma [14, 15]. The nasopharyngeal LEC patients were predominantly male, with a male to female ratio of 2.73:1. Moreover, our data showed that the incidence of nasopharyngeal LEC is much higher within the nonwhite/black population than in the white population. This is attributed to the prevalence and distribution of cancer-related viruses, such as EBV and HPV[16, 17]. Nasopharyngeal LEC patients can remain asymptomatic for a long time because nasopharynx is a clinically occult site. Nonspecific symptoms often caused delay a definitive diagnosis, one case series reported a mean delay period of 7.2 months[18]. Consequently, more than 90% of patients with a diagnosis of nasopharyngeal LEC present locally or regionally advanced disease [19]. Additionally, the previous studies provided those patients had a high incidence of distant metastases, ranging from 20% to 40%[20, 21]. In our study, 91.5% of nasopharyngeal LEC patients had regional or distant metastases, and 81.0% of cases had lymph node metastases. Due to anatomical limitations on surgical interventions, radiotherapy is undoubtedly the preferred choice of treatment and chemotherapy is combined in advanced disease[21]. For example, in the present study, only 20.8% of nasopharyngeal LEC patients underwent surgery, whereas 72.8% patients received chemotherapy, and 90.6% patients received radiotherapy.

Compared with nasopharyngeal LEC, non-nasopharyngeal LEC is much rarer. Until now, only three large case series have been reported. Ma et al. reported a cohort of 69 salivary gland LEC patients in China[7], while Dubey et al. reported 34 non-nasopharyngeal LEC patients in the United States between 1950 and 1994[22]. In 2016, Chan JY et al. reported 378 patients with non-nasopharyngeal LEC from the SEER database between 1973-2011[23]. All of these studies suggested that non-nasopharyngeal LECs were often located in the oropharynx, salivary gland, tonsil, tongue, etc. Chan JY and Dubey reported that the primary occurrence was in an oropharyngeal site in the majority of cases, followed by the salivary gland, while the present study found that the salivary gland was the most common site of non-nasopharyngeal LEC[23]. One possible reason behind this discrepancy is the difference in time span. Chan JY and Dubey’s studies recruited patients from 1950 to 1994 and 1973-2011, respectively. Our study retrospectively analyzed LEC patients from 1988 to 2013. The other reason is the diagnosis of LEC. In Chan JYK’s study, they included all non-nasopharyngeal LEC patients, while our study identified patients with only primary non-nasopharyngeal LEC. In addition, the clinical characteristics of non-nasopharyngeal LEC patients from the present study are consistent with those of previous reports. Our study found that most non-nasopharyngeal patients were men, with a male to female ratio of 1.6:1. Most non-nasopharyngeal LEC patients were aged younger than 60 years (62.5%) and were white (72.7%). Lymph node metastases in non-nasopharyngeal LEC are common with the incidence ranged from 10% to 50%. In the present study, the incidence of lymph node metastases was 70.2%, which is much higher than that from previous studies. One possibility for this discrepancy is that the lymph node status of 125 out of the 256 (48.8%) non-nasopharyngeal LEC patients was unknown in the present study. In addition, most non-nasopharyngeal LEC patients had advanced-stage (III/IV) disease, ranging from 59.4% to 80.2% at diagnosis in these studies. Our data reported that 78.2% patients were diagnosed with an advanced stage. In our cohort, 186 patients were white; in contrast, 53 cases of advanced-stage disease occurred in patients who were nonwhite, including American Indian/AK Native, Asian/Pacific Islander. Our findings support those of Chan JY, who identified that 23.7% of their non-nasopharyngeal LEC patients were nonwhite/black.

A strong association of EBV with LEC has been reported in Southeast Asia, Greenland, and Alaska, but not in the white population, especially for salivary gland LEC[7, 24, 25]. Several previous studies from high incidence regions have observed EBV positivity in non-nasopharyngeal LEC cases; for example, Ma reported that all 38 Chinese salivary gland LEC with EBV encoded RNA positive[7]. One potential explanation is associated with the geographic distribution of the 2 major types of EBV. Type 1EBV is the most prevalent type worldwide, whearas EBV type 2 is only common in certain areas such as Alaska, where there is a much higher incidence of salivary gland LEC. Although the present study did not analyze the EBV status in LEC patients due to inadequate information from SEER database, the 5-year OS of non-nasopharyngeal LEC was 81.1%, which is relatively lower than 90% reported by Ma et al., thus suggesting a difference in etiology and a different relationship with EBV. Besides, previous studies did not support a positive relationship of EBV with other non-nasopharyngeal, non-salivary gland LECs. For example, Singhi et al. found that all 22 patients with oropharynx LEC were HPV-P16 positive rather than EBV [6]. Chow et al. also reported 5 patients with intraoral LEC who were EBV negative. Of course, this relationship required to be further validated in the future.

With the development of molecular biology, an increasing number of potential biomarkers have been identified for furthering LEC developments. Previous studies have demonstrated the molecular signature of LEC, which includes BCL-2 overexpression, low rates of EGFR mutation, absence of HER2 and p53 expression[23]. BCL-2 overexpression may also render LEC patients sensitive to treatment with chemotherapeutic agents that target the apoptotic pathways associated with bcl-2[26]. In our cohort, 684 out of 1205 LEC patients received chemotherapy, and the prognosis data showed that chemotherapy could prolong the overall survival of LEC patients, but without significantly statistical difference. Because 341 of the LEC patients did not have accurate chemotherapy information, the role of chemotherapy in LEC patients needs to be confirmed in a large case study. Because of the similarities between the histology of LEC and nasopharyngeal carcinoma, the chemotherapy regimens commonly used for NPC may also work in LEC patients. The low occurrence rate of EGFR mutations and Her2 expression limits the utility of EGFR-TKIs as well as anti-Her2 antibodies. However, patient who received those treatments often have a good prognosis[8]. Similar to NPC, LEC of the oral cavity and pharynx has been demonstrated to be sensitive to radiotherapy. In the present study, radiotherapy could significantly improve the prognosis of LEC patients, and radiotherapy could decrease the risk of death by 49%. Postoperative radiotherapy was recommended for indications including non-R0 resection, stage T4, and lymph node metastases. In our cohort, 16.2% and 77.6% of LEC patients were diagnosed with T4 stage and lymph node metastases, respectively. Radiation after surgery could significantly prolong the survival of LEC patients who underwent surgery. In addition, the prognostic and therapeutic importance of EBV positivity and PD-L1 expression in nasopharyngeal carcinoma requires an investigation on whether LEC can be a potential immunotherapy target, similar to nasopharyngeal carcinoma, which has a strong immune response. Theoretically, lymphocyte compounds should be observed in the LEC tumor environment, and immunotherapy through the inhibition of immune suppression, such as through PD-L1 inhibitors, has promising prospects for LEC treatment.

In accordance with other reports, we found that patients with LEC of the oral cavity and pharynx had a much better prognosis. Almost 50% of the LEC patients could survive for 20 years in our cohort. In addition, the survival analysis in both unmatched cohort and matched cohort did not yield any significant difference in survival time between nasopharyngeal LEC and non-nasopharyngeal LEC patients. Therefore, we constructed a prognostic nomogram for individualized survival prediction of LEC patients using the long-term follow-up data from the SEER database. With this easy-to-use scoring system, both physicians and patients could calculate the survival probability of individual LEC patients. When a prognostic nomogram is completed, the validation of the nomogram is essential to avoid overfitting the model and determine its generalizability[27].[28, 29]. For common cancers, validation of the prognostic nomogram should be performed in the primary cohort and an independent cohort. However, this prognostic nomogram for LEC patients can only be validated in the primary cohort due to the rarity of LEC. In addition, several important prognostic indices were not included in this prognostic nomogram such as serum tumor markers. Further studies with more comprehensive information are required to confirm the accuracy of this nomogram.

In the present study, we described the clinicopathological characteristics and survival of patients with LEC of the oral cavity and pharynx. The results showed that patients with LEC of the oral cavity and pharynx often had a favorable prognosis. Old age, lymph node and distant metastases, surgery and radiotherapy were significantly associated with prognosis. Non-nasopharynx LEC patients were more likely to be young and male and have an undifferentiated grade and late TNM stage, but no any significant difference in prognosis between nasopharyngeal LEC and non-nasopharyngeal LEC patients could be observed. Meanwhile, we also constructed the first prognostic nomogram to predict the individual survival. In conclusion, the present study is the largest series concerning on LEC of the oral cavity and pharynx, and these results are vital to disease management and future prospective studies for this rare cancer.

## Supporting information

S1 Table 1: Characteristics of nasopharynx LEC patients and non-nasopharynx LEC patients.

S2 Table 2: Characteristics of nasopharynx LEC patients and non-nasopharynx LEC patients after PSM analysis in matched cohort.

## Reference

1. Kim YJ, Hong HS, Jeong SH, Lee EH, Jung MJ. Lymphoepithelial carcinoma of the salivary glands. Medicine (Baltimore). 2017;96(7):e6115. Epub 2017/02/17. doi: 10.1097/MD.0000000000006115. PubMed PMID: 28207533; PubMed Central PMCID: PMCPMC5319522.

2. Zhan KY, Nicolli EA, Khaja SF, Day TA. Lymphoepithelial carcinoma of the major salivary glands: Predictors of survival in a non-endemic region. Oral Oncol. 2016;52:24–9. Epub 2015/11/09. doi: 10.1016/j.oraloncology.2015.10.019. PubMed PMID: 26547125.

3. Coskun BU, Cinar U, Sener BM, Dadas B. Lymphoepithelial carcinoma of the larynx. Auris Nasus Larynx. 2005;32(2):189–93. Epub 2005/05/27. doi: 10.1016/j.anl.2004.11.014. PubMed PMID: 15917178.

4. Schneider M, Rizzardi C. Lymphoepithelial carcinoma of the parotid glands and its relationship with benign lymphoepithelial lesions. Arch Pathol Lab Med. 2008;132(2):278–82. Epub 2008/02/07. doi: 10.1043/1543-2165(2008)132[278:LCOTPG]2.0.CO;2. PubMed PMID: 18251590.

5. Acuna G, Goma M, Temprana-Salvador J, Garcia-Bragado F, Alos L, Ordi J, et al. Human papillomavirus in laryngeal and hypopharyngeal lymphoepithelial carcinoma. Mod Pathol. 2018. Epub 2018/12/16. doi: 10.1038/s41379-018-0188-2. PubMed PMID: 30552415.

6. Singhi AD, Stelow EB, Mills SE, Westra WH. Lymphoepithelial-like carcinoma of the oropharynx: a morphologic variant of HPV-related head and neck carcinoma. Am J Surg Pathol. 2010;34(6):800–5. Epub 2010/04/28. doi: 10.1097/PAS.0b013e3181d9ba21. PubMed PMID: 20421782.

7. Ma H, Lin Y, Wang L, Rao H, Xu G, He Y, et al. Primary lymphoepithelioma-like carcinoma of salivary gland: sixty-nine cases with long-term follow-up. Head Neck. 2014;36(9):1305–12. Epub 2013/08/24. doi: 10.1002/hed.23450. PubMed PMID: 23966284.

8. Zhao W, Deng N, Gao X, Chen TB, Xie J, Li Q, et al. Primary lymphoepithelioma-like carcinoma of salivary glands: a clinicopathological study of 21 cases. Int J Clin Exp Pathol. 2014;7(11):7951–6. Epub 2015/01/01. PubMed PMID: 25550837; PubMed Central PMCID: PMCPMC4270609.

9. Ke SJ, Wang P, Xu B. Clear cell adenocarcinoma of the lung: a population-based study. Cancer Manag Res. 2019;11:1003–12. Epub 2019/02/19. doi: 10.2147/CMAR.S187370. PubMed PMID: 30774428; PubMed Central PMCID: PMCPMC6349083.

10. Qin BD, Jiao XD, Liu K, Wu Y, He X, Liu J, et al. Clinical, pathological and treatment factors associated with the survival of patients with primary pulmonary salivary gland-type tumors. Lung Cancer. 2018;126:174–81. Epub 2018/12/12. doi: 10.1016/j.lungcan.2018.11.010. PubMed PMID: 30527184.

11. Hammas N, Benmansour N, El Alami El Amine MN, Chbani L, El Fatemi H. Lymphoepithelial carcinoma: a case report of a rare tumor of the larynx. BMC Clin Pathol. 2017;17:24. Epub 2017/12/06. doi: 10.1186/s12907-017-0061-0. PubMed PMID: 29204101; PubMed Central PMCID: PMCPMC5702189.

12. Henke C, Rieger J, Hartmann S, Middendorp M, Steinmetz H, Ziemann U. Paraneoplastic cerebellar degeneration associated with lymphoepithelial carcinoma of the tonsil. BMC Neurol. 2013;13:147. Epub 2013/10/19. doi: 10.1186/1471-2377-13-147. PubMed PMID: 24134642; PubMed Central PMCID: PMCPMC4016266.

13. Ibrahimov M, Yilmaz M, Celal MH, Mamanov M, Yollu U, Ozek H. Lymphoepithelial carcinoma of the larynx. J Craniofac Surg. 2013;24(3):1049. Epub 2013/05/30. doi: 10.1097/SCS.0b013e3182700cd9. PubMed PMID: 23714948.

14. Ozyar E, Selek U, Laskar S, Uzel O, Anacak Y, Ben-Arush M, et al. Treatment results of 165 pediatric patients with non-metastatic nasopharyngeal carcinoma: a Rare Cancer Network study. Radiother Oncol. 2006;81(1):39–46. Epub 2006/09/13. doi: 10.1016/j.radonc.2006.08.019. PubMed PMID: 16965827.

15. Zubizarreta PA, D’Antonio G, Raslawski E, Gallo G, Preciado MV, Casak SJ, et al. Nasopharyngeal carcinoma in childhood and adolescence: a single-institution experience with combined therapy. Cancer. 2000;89(3):690–5. Epub 2000/08/10. PubMed PMID: 10931470.

16. Cardesa A, Nadal A. Carcinoma of the head and neck in the HPV era. Acta Dermatovenerol Alp Pannonica Adriat. 2011;20(3):161–73. Epub 2011/12/02. PubMed PMID: 22131117.

17. Wenig BM. Lymphoepithelial-like carcinomas of the head and neck. Semin Diagn Pathol. 2015;32(1):74–86. Epub 2015/03/26. doi: 10.1053/j.semdp.2014.12.004. PubMed PMID: 25804344.

18. Leong JL, Fong KW, Low WK. Factors contributing to delayed diagnosis in nasopharyngeal carcinoma. J Laryngol Otol. 1999;113(7):633–6. Epub 1999/12/22. PubMed PMID: 10605559.

19. Fandi A, Altun M, Azli N, Armand JP, Cvitkovic E. Nasopharyngeal cancer: epidemiology, staging, and treatment. Semin Oncol. 1994;21(3):382–97. Epub 1994/06/01. PubMed PMID: 8209270.

20. Ahmad A, Stefani S. Distant metastases of nasopharyngeal carcinoma: a study of 256 male patients. J Surg Oncol. 1986;33(3):194–7. Epub 1986/11/01. PubMed PMID: 3773537.

21. Wu FY, Yang ES, Willey CD, Ely K, Garrett G, Cmelak AJ. Refractory lympho-epithelial carcinoma of the nasopharynx: a case report illustrating a protracted clinical course. Head Neck Oncol. 2009;1:18. Epub 2009/06/17. doi: 10.1186/1758-3284-1-18. PubMed PMID: 19527509; PubMed Central PMCID: PMCPMC2704192.

22. Dubey P, Ha CS, Ang KK, El-Naggar AK, Knapp C, Byers RM, et al. Nonnasopharyngeal lymphoepithelioma of the head and neck. Cancer. 1998;82(8):1556–62. Epub 1998/04/29. PubMed PMID: 9554534.

23. Chan JY, Wong EW, Ng SK, Vlantis AC. Non-nasopharyngeal head and neck lymphoepithelioma-like carcinoma in the United States: A population-based study. Head Neck. 2016;38 Suppl 1:E1294–300. Epub 2015/09/01. doi: 10.1002/hed.24215. PubMed PMID: 26316257.

24. Hamilton-Dutoit SJ, Therkildsen MH, Neilsen NH, Jensen H, Hansen JP, Pallesen G. Undifferentiated carcinoma of the salivary gland in Greenlandic Eskimos: demonstration of Epstein-Barr virus DNA by in situ nucleic acid hybridization. Hum Pathol. 1991;22(8):811–5. Epub 1991/08/01. PubMed PMID: 1651284.

25. Saku T, Cheng J, Jen KY, Tokunaga M, Li J, Zhang W, et al. Epstein-Barr virus infected lymphoepithelial carcinomas of the salivary gland in the Russia-Asia area: a clinicopathologic study of 160 cases. Arkh Patol. 2003;65(2):35–9. Epub 2004/09/11. PubMed PMID: 15357246.

26. Kang MH, Reynolds CP. Bcl-2 inhibitors: targeting mitochondrial apoptotic pathways in cancer therapy. Clin Cancer Res. 2009;15(4):1126–32. Epub 2009/02/21. doi: 10.1158/1078-0432.CCR-08-0144. PubMed PMID: 19228717; PubMed Central PMCID: PMCPMC3182268.

27. Liang W, Zhang L, Jiang G, Wang Q, Liu L, Liu D, et al. Development and validation of a nomogram for predicting survival in patients with resected non-small-cell lung cancer. J Clin Oncol. 2015;33(8):861–9. doi: 10.1200/JCO.2014.56.6661. PubMed PMID: 25624438.

28. Tang LQ, Li CF, Li J, Chen WH, Chen QY, Yuan LX, et al. Establishment and Validation of Prognostic Nomograms for Endemic Nasopharyngeal Carcinoma. J Natl Cancer Inst. 2016;108(1). doi: 10.1093/jnci/djv291. PubMed PMID: 26467665.

29. Won YW, Joo J, Yun T, Lee GK, Han JY, Kim HT, et al. A nomogram to predict brain metastasis as the first relapse in curatively resected non-small cell lung cancer patients. Lung Cancer. 2015;88(2):201–7. doi: 10.1016/j.lungcan.2015.02.006. PubMed PMID: 25726044.

